# Cell engulfment defines spatially distinct competitive metabolic niches associated with clinical outcomes in colorectal cancer

**DOI:** 10.1101/2025.07.15.662825

**Authors:** Emir Bozkurt, Batuhan Kisakol, Heiko Dussmann, Federico Lucantoni, Anna Sturrock, Mohammadreza Azimi, Sanghee Cho, Elizabeth McDonough, Joana Fay, Tony O’Grady, John P. Burke, Niamh McCawley, Deborah A. McNamara, Daniel B. Longley, Fiona Ginty, Jochen H.M. Prehn

**Affiliations:** Department of Physiology and Medical Physics & RCSI Centre for Systems Medicine, RCSI University of Medicine and Health Sciences, 123 St Stephen’s Green, Dublin 2, Ireland; Department of Mechanical Engineering, Virginia Tech, Blacksburg, VA, 24061, U.S.A.; Laboratory of Cellular Stress and Cell Death Pathways, Centro de Investigación Príncipe Felipe (CIPF), Valencia, Spain; Institute of Microbiology and Virology, Riga Stradins University, LV-1067 Riga, Latvia; GE HealthCare Technology & Innovation Center (Formerly GE Research Center), 1 Research Circle, Niskayuna, NY, 12309, U.S.A.; Department of Pathology, RCSI University of Medicine and Health Sciences, Beaumont Hospital, Dublin 9, Ireland; Department of Surgery, RCSI University of Medicine and Health Sciences, Beaumont Hospital, Dublin 9, Ireland; Queens University, School of Medicine, Dentistry and Biomedical Sciences, Patrick G Johnston Centre for Cancer Research, 97 Lisburn Road, Belfast, BT9 AE, U.K.

**Keywords:** Cell competition, cell-in-cell, entosis, emperipolesis, cell engulfment, cancer stem cell, T cell, colorectal cancer

## Abstract

Cell competition is an emerging mechanism in which mammalian tissues maintain homeostasis by eliminating less fit (loser) cells through direct interactions with fitter (winner) neighbouring cells. In cancer, these competitive interactions may drive tumour evolution; however, spatial organisation and clinical relevance of these events remain poorly understood. One mechanism by which winner cells eliminate loser cells is engulfment, resulting in cell-in-cell (CIC) formation. Although CICs have been observed in many tumour types for over a century, their cellular composition, spatial context, interactions with the tumour microenvironment, and biological significance in human cancers remain unclear.

Here, we systematically characterised the cellular identity and functional states of CICs *in situ*, examined their spatial interactions within the tumour microenvironment, and assessed their clinical relevance using spatially resolved single-cell data from a large cohort of colorectal cancer patients. We demonstrate that CICs occur predominantly between cancer cells but also involve cancer stem cell (CSC)-like populations and cytotoxic T cells. Engulfed (inner) cancer and CSC-like cells display molecular features consistent with a loser-cell phenotype, including increased apoptosis and reduced proliferation, whereas outer cancer cells exhibit winner-cell features such as upregulated glycolysis. Live-cell time-lapse experiments demonstrate that glucose accumulates in inner cells during lysosomal degradation following cell engulfment. Spatial analysis further revealed distinct CIC neighbourhoods, which we defined based on proximity to engulfment events. Cells within these regions, particularly CSC-like cells and cytotoxic T cells, exhibit increased metabolic stress, suggesting local competition for nutrients. Importantly, the presence of cytotoxic T cells within CIC neighbourhoods and spatial co-occurrence patterns between cancer cells and CSC-like populations are associated with improved patient outcomes. Together, our findings demonstrate that cell engulfment defines spatially organised competitive niches and may reflect cell competition within complex tumour microenvironments.

## Introduction

Cell competition is an evolutionarily conserved process through which tissues regulate architecture and maintain homeostasis. It is a close-proximity interaction in which cells compare their fitness through direct contact and eliminate neighbours with lower fitness^1,2^. Elimination typically involves activation of apoptosis in loser cells followed by their engulfment by neighbouring cells^1,3–5^. Intriguingly, loser cells have also been observed to remain viable, without any evidence of apoptosis, while residing inside winner cells^6^. In normal tissues, including the intestinal epithelium, cell competition functions as a pluripotency surveillance mechanism that eliminates slowly proliferating stem cells and shapes mammalian tissue morphogenesis by overcoming cellular heterogeneity^2,7^. In colorectal cancer (CRC), malignant cells may evade cell competition until oncogenic mutations such as *KRAS, TP53, MYC*, and *EGFR* increase their cellular fitness and confer winner-cell status^2,8–10^. Winner-cell status is also associated with metabolic rewiring, particularly increased glucose uptake and lactate production^11,12^, and can be further modified by alterations in JNK signalling^4,9^ and BCL-2 family proteins^5,6,13,14^. Whether cancer cells exploit competitive interactions to their advantage, where in the tumour these events occur, which cell types participate, and how these processes relate to clinical outcomes remain poorly understood.

Engulfment of neighbouring cells by cancer cells has been documented in tumours for over a century as cell-in-cell (CIC) structures^15^. CICs occur more frequently in tumours than matched normal tissues^16^, and have been studied as a mechanism of cell competition through entosis^17–19^. Studies suggest that oncogenic mutations such as *KRAS*^18^, *RAC*^18^, *TP53*^20,21^, *MYC*^22^, and *EGFR*^20^ promote winner-cell status, enabling cancer cells to engulf and eliminate neighbouring cells. Although the underlying mechanisms are unclear, *in vitro* studies show that CICs are associated with increased expression of glycolytic genes^23^, while AMPK^24^ and JNK^25^ signalling can alter winner/loser cell status. Perturbations in BCL-2 family proteins can also promote survival and escape of engulfed inner cells^16,17^. In addition to genetic changes, metabolic^24^, apoptotic^25–27^, and proliferative^28^ stress increase the frequency of cell engulfment events and confer survival advantages to outer cells. Conversely, during interactions between tumour-reactive T cells and cancer cells, outer cancer cells may be eliminated by T-cell-mediated cytotoxicity, whereas inner cells are protected from both direct contact with reactive T cells and exposure to lytic granules released by immune cells^29^. Cancer cells can also engulf and eliminate nearby T cells and NK cells through emperipolesis^30–32^ and enclysis^33^. Together, these findings suggest that cancer cells may exploit engulfment through diverse, context-dependent interactions that remodel the tumour microenvironment and promote immune evasion.

Although CICs are commonly observed in cancers, their clinical significance varies across tumour types. In head and neck^34,35^, pancreatic^36^, oral^37^, hepatocellular^38^, and lung cancers^20^, CICs are associated with poor prognosis, whereas in esophageal^39^ and breast cancers^40,41^, CICs correlate with improved clinical outcomes. Furthermore, tumour regions in which CICs occur and cell types involved in these events have shown inconsistent clinical associations^27,40,42^. These discrepancies suggest that the functional consequences of CICs may depend on tumour context, cellular composition, or the local microenvironment in which these interactions occur. However, the diversity of CICs within the tumour microenvironment, including the identity, spatial distribution, and functional state of the participating cells, and their tissue-level and clinical significance in human cancers remain poorly understood.

Here, we combined multiplex single-cell imaging with spatial analysis to characterise cell engulfment events *in situ* and to determine their cellular identity, functional states, spatial interactions with the tumour microenvironment, and clinical relevance in colorectal tumours. We showed that CICs predominantly involve cancer cells, with smaller proportions involving cancer stem cell-like (CSC-like) cells, cytotoxic T cells, and monocytes. Tumour regions containing CICs were associated with reduced cytotoxic T cell levels. We described protein expression profiles related to metabolism, apoptosis, and proliferation in inner and outer cells, and showed that glucose accumulated in inner cells during lysosomal degradation following cell engulfment. CICs were increased in tumours exhibiting lymphovascular or extramural venous invasion. Finally, we defined “CIC neighbourhoods” as spatially distinct intratumoural niches consistent with cell competition and demonstrated their clinical relevance in CRC.

## Results

### CICs predominantly occur between cancer cells and are associated with reduced cytotoxic T cell levels

We performed Cell DIVE multiplex imaging to profile single-cell expression of 60 proteins (Table S1) in 344 regions of tumour cores from 124 colorectal cancer patients. Cells were classified based on marker expression into cancer, stromal, endothelial, macrophage, monocyte, cytotoxic T, helper T, regulatory T, and cancer stem cell–like (CSC-like) populations (Figure 1A). Next, we manually annotated cell-in-cell structures (CICs) in HALO software if they exhibited evidence of a complete internalisation of an entire cell within another cell and if the event met at least five of the six following criteria: (1) nucleus of inner cell, (2) cytoplasm of inner cell, (3) membrane of inner cell, (4) nucleus of outer cell, (5) cytoplasm of outer cell, and (6) visible vacuolar space between inner and outer cells (Figure 1A,B, Figure S1), consistent with previous studies^16,20,43^. In total, 443 CICs were identified using virtual H&E images together with nuclear (DAPI), cytoplasmic (AE1, PCK26, EPCAM), and membrane (NAK, PCAD) markers.

**Figure 1:**
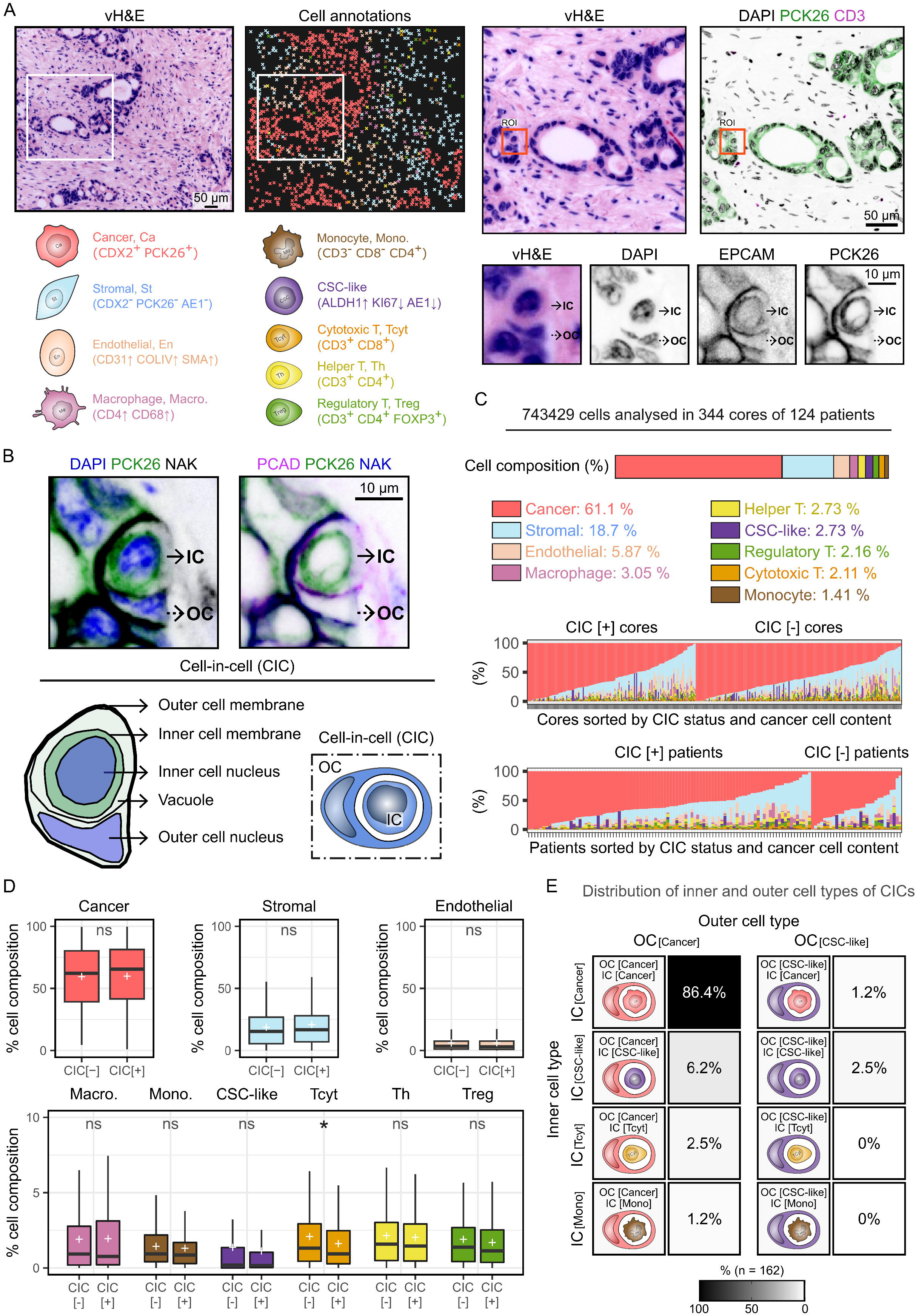
Characterisation of cell-in-cell (CIC) events and their association with cellular composition of colorectal tumours. **A**. Classification of cancer cells, stromal cells, endothelial cells, macrophages, monocytes, CSC-like cells, cytotoxic T cells, helper T cells, and regulatory T cells within CRC tumours. The region of interest (right) depicts a smaller area showing a CIC event between two cancer cells (red). **B**. Characterisation of CIC events in CRC tissue sections. Combination of vH&E, nuclear (DAPI), cytoplasmic (PCK26, AE1) and membrane (PCAD, NAK) markers was used to detect CICs (IC: inner cell, OC: outer cell). A schematic representing the characteristics of a CIC was generated based on the staining. **C**. Cellular composition of CRC tumours based on cell types classified in this study. In total, 743429 cells were analysed across 344 cores from 124 patients. Distribution of cellular composition in CIC positive (CIC [+]) and CIC negative (CIC [-]) cores and patients is shown. **D**. Comparison of cancer, stromal, endothelial, macrophage, monocyte, CSC-like cell, cytotoxic T, helper T, and regulatory T cell compositions between CIC [-] (n = 189) and CIC [+] (n = 155) cores. Groups were compared using unpaired t-test, data are presented as median, interquartile range, and mean (+). *P < 0.05, **P < 0.01, ns: not significant. **E**. Distribution of CICs (n = 162) by inner cell (IC) and outer cell (OC) types. Shades of grey indicate proportion of each group (darker colour represents higher value).

Among 743,429 cells analysed, cancer cells comprised 61.1% of the population, followed by stromal (18.7%), endothelial (5.87%), macrophages (3.05%), helper T cells (2.73%), CSC-like cells (2.73%), regulatory T cells (2.16%), cytotoxic T cells (2.11%), and monocytes (1.41%) (Figure 1C). Cell composition varied substantially across tumour cores, indicating high intra- and inter-tumour heterogeneity within the cohort.

Comparison of CIC-positive (CIC > 0) and CIC-negative (CIC = 0) CRC cores revealed significantly lower proportions of cytotoxic T cells in CIC-positive regions, whereas other cell populations did not differ significantly (Figure 1D). These results suggest that CIC formation is associated with tumour regions exhibiting reduced cytotoxic T cell infiltration.

We next characterised the cellular composition of inner cells (IC) and outer cells (OC) of CICs. All CICs contained cancer or CSC-like cells as outer cells. The majority of CICs (86.4%) occurred between cancer cells. Among outer cancer cells, 6.2% engulfed CSC-like cells, 2.5% engulfed cytotoxic T cells, and 1.2% engulfed monocytes. CSC-like outer cells accounted for 3.7% of CICs and engulfed either cancer cells (1.2%) or CSC-like cells (2.5%) (Figure 1E).

Together, these findings indicate that CIC formation in colorectal cancer predominantly occurs through homotypic cancer cell interactions, while heterotypic events involving immune or CSC-like cells are relatively rare.

### Inner and outer cells exhibit metabolic and apoptotic profiles consistent with cell competition

We compared the single-cell expression profiles of proteins involved in apoptosis, metabolism, and proliferation between CICs and non-engulfed cells (Figure 2, Figure S2). Inner and outer cancer cells were compared with non-engulfed cancer cells, and a similar approach was used for CSC-like cells. Inner cytotoxic T cells were also compared with non-internalised cytotoxic T cells to examine changes associated with emperipolesis^15,30^.

**Figure 2:**
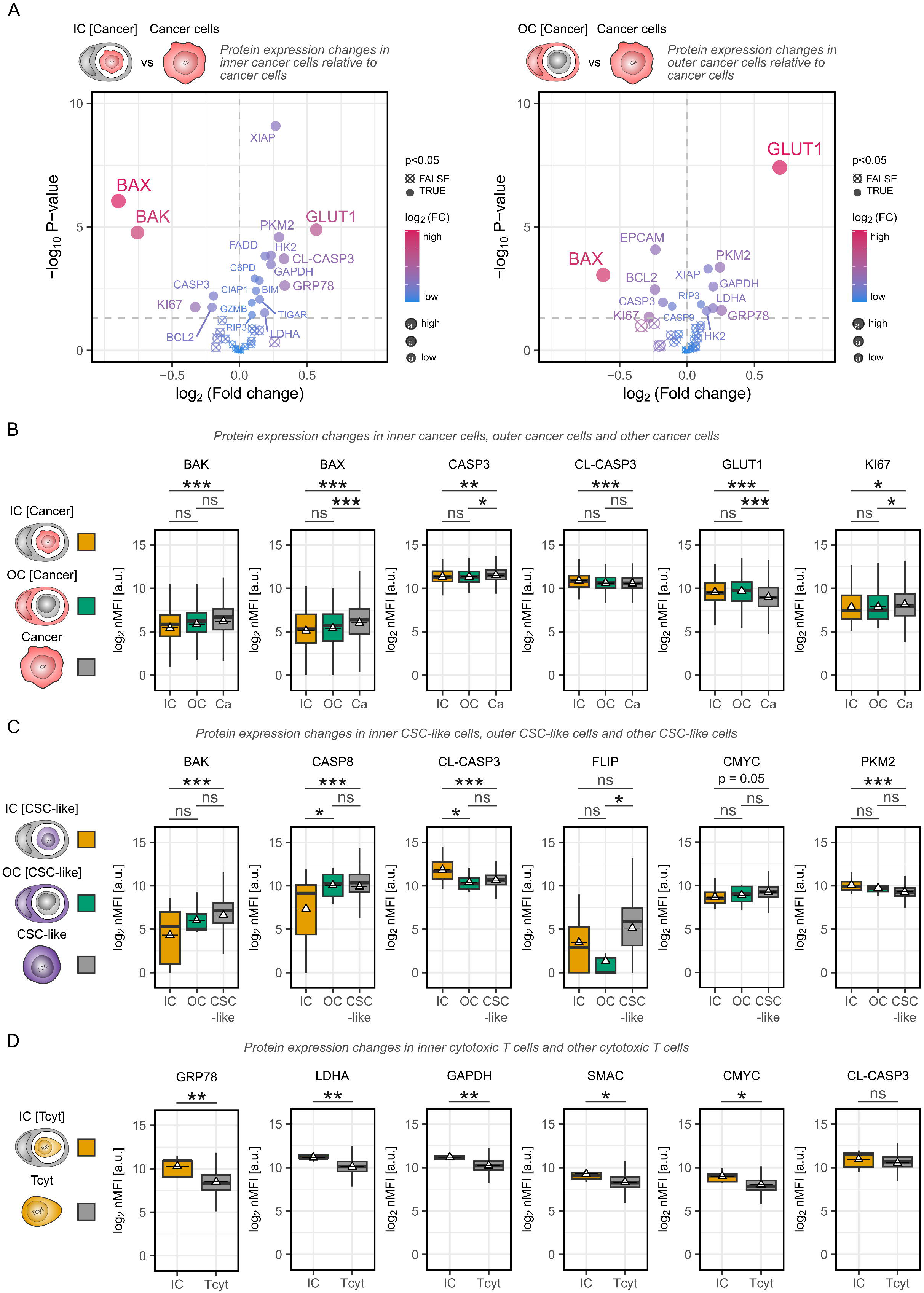
Metabolic, apoptotic, and proliferation profiles of inner and outer cells. **A**. Volcano plots showing differential protein expression between inner cancer cells (IC [cancer], n = 171, left) and outer cancer cells (OC [cancer], n = 179, right) compared to other cancer cells (Ca, n = 444725). Proteins with significant changes are indicated as filled circles. The dashed horizontal line denotes P = 0.05. Colour and point size represent log_2_ fold change. **B**. Comparison of log_2_ normalised mean fluorescence intensity (nMFI) values of selected differentially expressed proteins in IC [cancer] (n = 171), OC [cancer] (n = 179), and other cancer cells (Ca, n = 444725). Triangles indicate group means. Groups were compared using one-way ANOVA followed by Tukey’s HSD post-hoc test for multiple comparisons. *P < 0.05, **P < 0.01, ***P < 0.001, ns: not significant. **C**. Comparison of log_2_ nMFI values of differentially expressed proteins in IC [CSC-like] (n = 13), OC [CSC-like] (n = 6), and other CSC-like cells (n = 20255). Triangles indicate group means. Groups were compared using one-way ANOVA followed by Tukey’s HSD post-hoc test for multiple comparisons. *P < 0.05, **P < 0.01, ***P < 0.001, ns: not significant. **D**. Comparison of log_2_ nMFI values of differentially expressed proteins between IC [Tcyt] (n = 5) and other cytotoxic T cells (Tcyt, n = 15668). Triangles indicate group means. Groups were compared using an unpaired t-test. *P < 0.05, **P < 0.01, ***P < 0.001, ns: not significant.

Inner cancer cells exhibited significantly reduced expression of mitochondrial apoptosis regulators, with BAX and BAK showing the most pronounced decreases relative to other cancer cells. CASP3 and BCL2 were slightly reduced, whereas cleaved CASP3 (CL-CASP3) was elevated. Outer cancer cells showed similar but less pronounced changes and did not display increased CL-CASP3. Both inner and outer cancer cells exhibited elevated expression of metabolic and stress-associated proteins, including GLUT1, GRP78, PKM2, HK2, GAPDH, LDHA, and TIGAR. GLUT1 expression was slightly higher in outer cells than inner cells, although this difference was not statistically significant. Proliferation marker KI67 was reduced in both inner and outer cancer cells (Figure 2A,B).

Approximately 10% of CICs involved CSC-like cells (Figure 1E). Inner CSC-like cells displayed reduced BAK and CASP8 and increased CL-CASP3 and PKM2 compared with non-engulfed CSC-like cells. Outer CSC-like cells showed reduced FLIP expression. Inner CSC-like cells exhibited lower CASP8 and CMYC and higher CL-CASP3, consistent with loser-cell phenotype (Figure 2C).

A small proportion of CICs contained cytotoxic T cells as inner cells (2.5%) (Figure 1E). These cells exhibited increased expression of GRP78, LDHA, GAPDH, SMAC, and CMYC compared with non-engulfed cytotoxic T cells, whereas CL-CASP3 levels were unchanged (Figure 2D).

Together, these results indicate that CIC-associated cancer and CSC-like cells display distinct apoptotic and metabolic states consistent with cell competition. Inner cells exhibited elevated CL-CASP3, consistent with loser-cell phenotype, whereas outer cancer cells showed higher GLUT1 expression, suggesting a metabolically fitter, winner-cell phenotype. CIC formation was therefore associated with apoptotic remodelling, altered glucose metabolism, and reduced proliferation in tumour cells, while inner cytotoxic T cells primarily displayed metabolic and ER stress without clear evidence of apoptosis.

### CICs exhibit increased glucose uptake during lysosomal degradation of inner cells

To investigate glucose metabolism during CIC formation, we monitored uptake of the fluorescent glucose analogue 2-NBDG using live-cell time-lapse confocal microscopy in HCT116 colon cancer cells. Previous studies have shown that following CIC formation, most inner cells undergo lysosomal degradation within outer cells^16,17,32,44^. Consistent with this, 2-NBDG accumulated specifically in inner cells undergoing lysosomal degradation (IC LT[+]), whereas during early CIC stages prior to Lysotracker accumulation (IC LT[-]) 2-NBDG levels remained similar to neighbouring cells (Figure 3A,B). Quantification of 2-NBDG uptake revealed an approximately four-fold higher glucose uptake rate in degrading inner cells compared with neighbouring cancer cells (Figure 3C).

**Figure 3:**
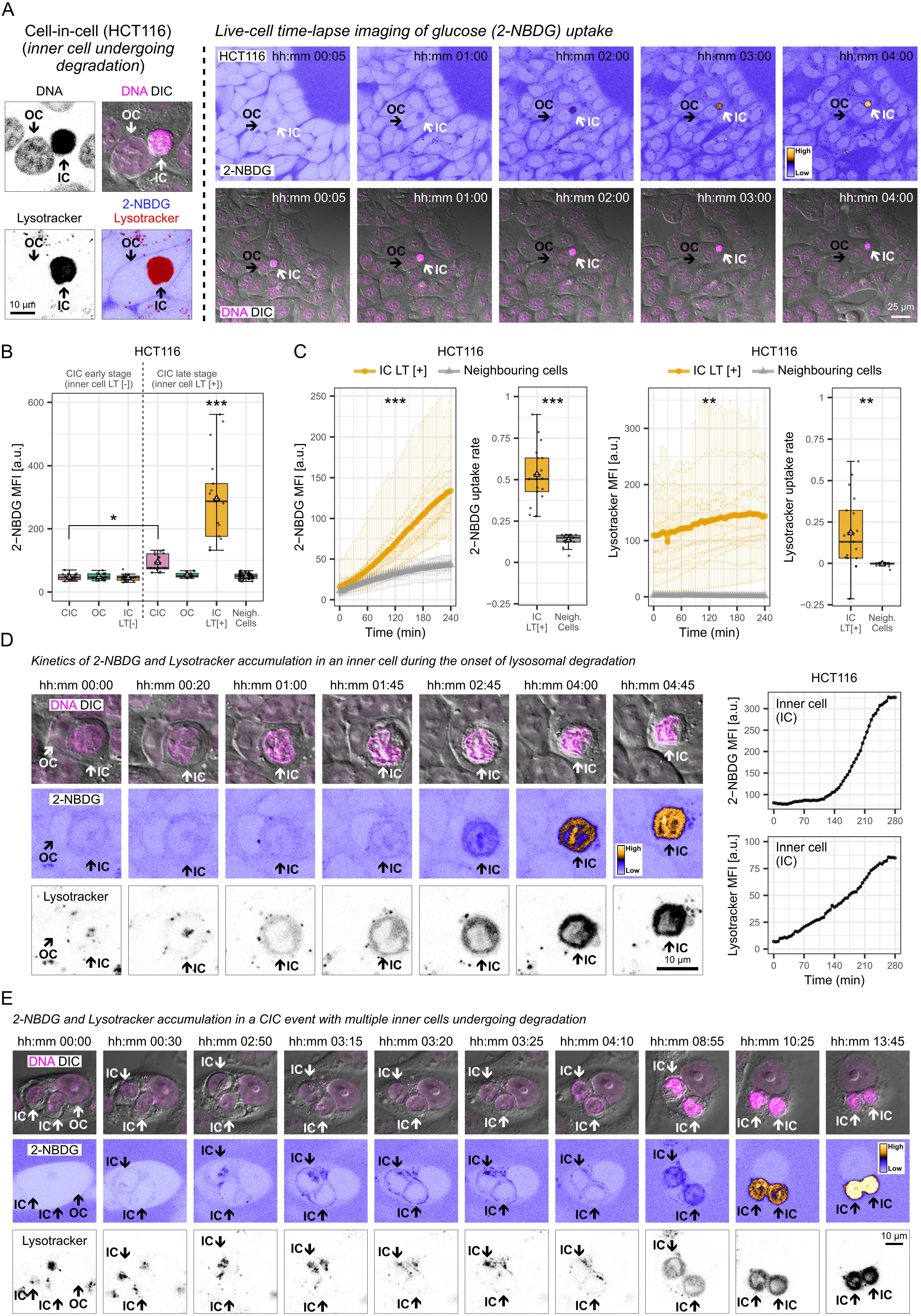
Glucose accumulates in inner cells during lysosomal degradation following cell engulfment. **A**. Representative confocal microscopy images of Hoechst (DNA), Lysotracker, and 2-NBDG (fluorescent glucose analogue) showing a late-stage inner cell undergoing degradation within an outer cell in HCT116 cells (left). Representative live-cell time-lapse confocal microscopy images showing accumulation of 2-NBDG in a field of view containing a CIC event with a late-stage inner cell. IC: inner cell, OC: outer cell. **B**. Quantification of 2-NBDG mean fluorescence intensity (MFI) in early-stage CICs, late-stage CICs, and neighbouring cells in HCT116. Early-stage CICs contain inner cells without Lysotracker accumulation (LT [-]), whereas late-stage CICs contain inner cells undergoing lysosomal degradation (LT [+]). Numbers of analysed cells for each group were: nCIC_ICLT[-] = 18, nOC_ICLT[-] = 18, nICLT[-] = 19, nCIC_ICLT[+] = 15, nOC_ICLT[+] = 15, nICLT[+] = 17, Neigh. cells = 72. Groups were compared using one-way ANOVA followed by Tukey’s HSD post-hoc test for multiple comparisons. *P < 0.05, ***P < 0.001. **C**. Comparison of 2-NBDG and Lysotracker accumulation kinetics in inner cells undergoing degradation within an outer cell (ICLT[+], n = 17) and neighbouring HCT116 cells (n = 16). 2-NBDG and Lysotracker uptake rates were calculated from the slopes of time-lapse measurements. Groups were compared using an unpaired t-test. **P < 0.01, ***P < 0.001. **D**. Representative live-cell time-lapse confocal microscopy images of DNA, Lysotracker, and 2-NBDG at the onset of lysosomal degradation in an inner cell. IC: inner cell, OC: outer cell. Quantification of 2-NBDG and Lysotracker MFI for the corresponding IC is shown. **E**. Representative live-cell time-lapse confocal microscopy images of DNA, Lysotracker, and 2-NBDG in multiple inner cells undergoing lysosomal degradation in a CIC event. IC: inner cell, OC: outer cell.

The onset of 2-NBDG accumulation partially coincided with Lysotracker signal. During early degradation, 2-NBDG was distributed throughout the inner cell, while Lysotracker signal was enriched at the cell periphery. Some Lysotracker-positive vesicles were also positive for 2-NBDG, indicating partial co-localisation (Figure 3D). In CICs containing multiple inner cells, 2-NBDG accumulated in each inner cell undergoing lysosomal degradation (Figure 3E). Inner cells progressively shrank and were ultimately degraded by the outer cells.

To assess glucose uptake in heterotypic CICs, we co-cultured HCT116 cells with Jurkat T lymphocytes and tracked LT[+] inner Jurkat cells within outer cancer cells. Similar to homotypic CICs, 2-NBDG accumulated at significantly higher levels in inner T cells undergoing lysosomal degradation (Figure S3).

We next examined GLUT1 expression in HCT116 cells by immunofluorescence. Consistent with live-cell imaging results, inner cells undergoing lysosomal degradation exhibited higher GLUT1 levels compared to neighbouring cells. This increase was evident even in CICs where only one inner cell was Lysotracker[+] while another remained Lysotracker[-]. CICs and outer cells containing LT[+] inner cells showed slightly elevated GLUT1 expression, although this difference was not statistically significant (Figure S3).

Together, these results indicate that initiation of inner cell degradation coincides with a rapid increase in glucose uptake within inner cells in both homotypic and heterotypic CICs. These metabolic changes appear transient and are consistent with previously reported rapid AMPK activation in loser (inner) cells during entosis^24^. Moreover, outer (winner) cells may acquire nutrients from the degraded inner (loser) cells.

### CICs are associated with invasive and mesenchymal clinicopathological features in CRC

We examined associations between CICs and clinicopathological, molecular, and clinical features in a cohort of stage I-III CRC patients. Patient demographics and tumour characteristics are summarised in Table S2 and Figure 4A. CIC distribution across tumours was highly heterogeneous and varied substantially between patients (Figure 4A). Overall, CICs were detected in 112 of 148 patients (75.6%), consistent with our previous observations in an independent CRC cohort (121/196, 61.7%)^16^.

**Figure 4:**
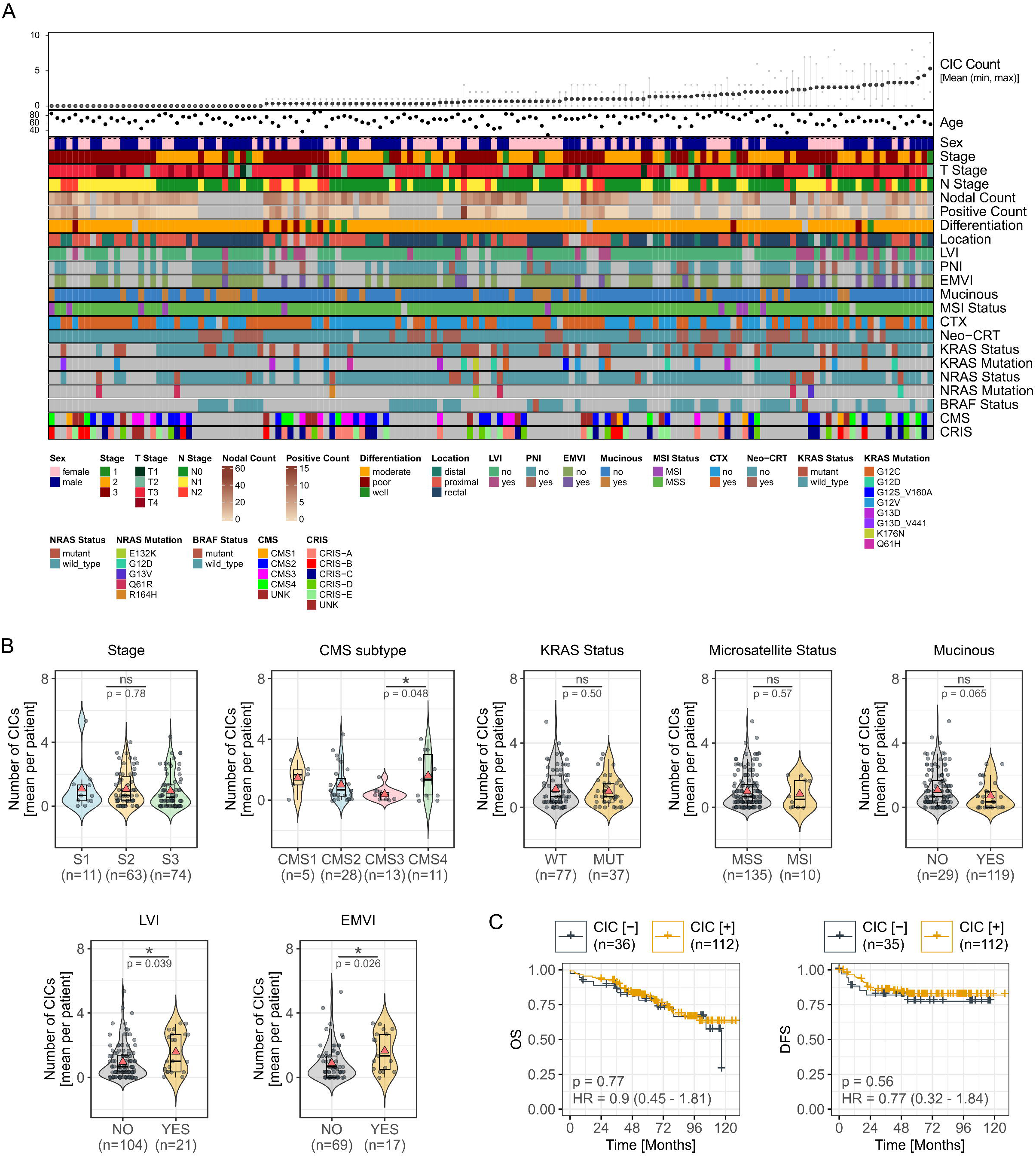
Association of CICs with clinicopathological and molecular features in colorectal cancer. **A**. Inter- and intra-patient heterogeneity in CIC counts (top) and overview of clinical, pathological, and molecular characteristics of the colorectal cancer cohort. Each column represents a patient and each row indicates a clinical or molecular feature (colour-coded). Patients are ordered along the x-axis by mean CIC counts (low to high). Missing data are shown in grey. **B**. Comparison of CIC counts by tumour stage, CMS subtype, KRAS status, microsatellite status, mucinous features, lymphovascular invasion, and extramural vascular invasion in CRC patients. Groups were compared using one-way ANOVA followed by Tukey’s HSD post-hoc test for multiple comparisons or by unpaired t-test. *P < 0.05, ns: not significant. **C**. Kaplan–Meier estimates of overall survival (OS) and disease-free survival (DFS) comparing the absence (CIC [-]) and presence (CIC [+]) of CICs in CRC patients. P values and hazard ratios (HR) from adjusted Cox proportional hazard models are shown.

In line with our previous study^16^, CIC presence was not associated with age, sex, tumour stage, nodal count, differentiation, tumour location, chemotherapy status, MSI status, mutational profile, or molecular subtype. However, CICs were significantly more frequent in tumours exhibiting lymphovascular invasion (LVI) and extramural venous invasion (EMVI), and were enriched in CMS4 compared to CMS3 tumours. A trend toward reduced CIC numbers was observed in tumours with mucinous features (p = 0.065) (Figure 4B). We next assessed the association between CIC presence and clinical outcomes using Kaplan–Meier estimates and Cox proportional hazard models adjusted for age and sex. Consistent with our previous CRC cohort^16^, CIC presence (CIC > 0) was not associated with overall survival (OS) [HR = 0.9, 95% CI: 0.45-1.81, p = 0.77] or disease-free survival (DFS) [HR = 0.77, 95% CI: 0.32-1.84, p = 0.56] (Figure 4C). Together, these results indicate that while CIC presence is not associated with survival outcomes, it is correlated with pathological features linked to tumour invasion and mesenchymal differentiation.

### CIC neighbourhoods define spatial niches enriched in CSC-like cells and are associated with metabolic adaptations and T cell-dependent survival

To examine whether CICs occupy spatially distinct tumour microenvironments, we analysed the composition of cells located within 20 *µ*m of the centre of a CIC, defined as the cell-in-cell (CIC) neighbourhood (Figure 5A). CIC neighbourhoods consisted predominantly of cancer cells (84.0%), followed by stromal cells (6.44%), CSC-like cells (4.12%), endothelial cells (1.63%), macrophages (1.46%), cytotoxic T cells (0.95%), regulatory T cells (0.69%), monocytes (0.51%), and helper T cells (0.17%). Cellular composition of CIC neighbourhoods varied across tumour cores and patients (Figure 5A).

**Figure 5:**
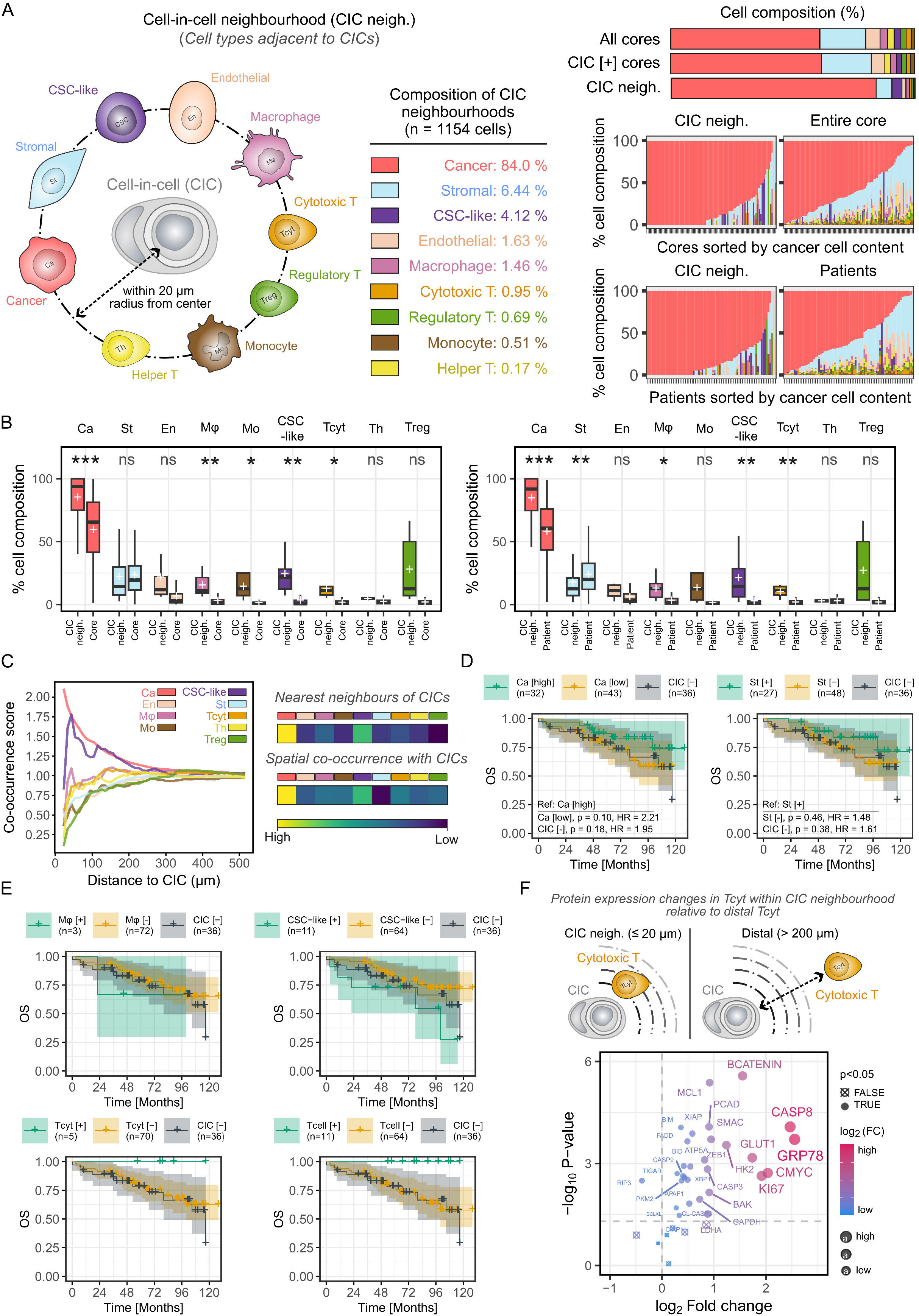
Spatial composition of cell-in-cell neighbourhoods and its association with patient survival. **A**. The term “cell-in-cell neighbourhood (CIC neigh.)” is used to define cell types that are in close proximity to a CIC (left). Cellular composition of CIC neighbourhoods based on cell types classified in this study (middle). In total, 1154 cells were analysed. Distribution of cellular composition in CIC neighbourhoods, entire cores (n = 155) and patients (n = 94) are shown (right). **B**. Pairwise comparisons of cancer, stromal, endothelial, macrophage, monocyte, CSC-like cell, cytotoxic T, helper T, and regulatory T cell compositions between CIC neighbourhoods and entire cores or patients. Groups were compared using paired t-test, data presented as the median, interquartile range, and mean (+). *P < 0.05, **P < 0.01, ***P < 0.001, ns: not significant. **C**. Co-occurrence probability and nearest-neighbour enrichment scores were calculated to analyse the spatial relationships of cell types within CIC neighbourhoods. Co-occurrence score estimates the likelihood that two cell types co-exist within the same spatial region, while nearest-neighbour enrichment score represents the likelihood that two cell types are immediate neighbours. Co-occurrence scores were plotted for each cell type as a function of distance from a CIC event. **D**. Kaplan–Meier estimates comparing overall survival in CRC patients based on the cancer (left) and stromal (right) composition of CIC neighbourhoods, as well as the absence of CICs (CIC[-]). Cancer cells were compared as high (Ca[high]) vs low (Ca[low]) abundance while stromal cells were compared by presence (St[+]) or absence (St[-]). P values and hazard ratios (HR) from Cox proportional hazard models adjusted for age and sex are shown. **E**. Kaplan–Meier estimates comparing overall survival in CRC patients based on the presence ([+]) and absence ([-]) of macrophages (Mφ), CSC-like cells (CSC-like), cytotoxic T cells (Tcyt), and T cells (T cell), as well as the absence of CICs (CIC[-]). **F**. Volcano plot showing differential protein expression between cytotoxic T cells (Tcyt) within CIC neighbourhood (n = 11) and distal Tcyts (n = 2567). Proteins with significant changes are indicated as filled circles. The dashed horizontal line denotes P = 0.05. Colour and point size represent log_2_ fold change. Groups were compared using an unpaired t-test.

Pairwise comparisons between CIC neighbourhoods and overall tumour composition within the same cores and patients revealed significantly higher proportions of cancer cells, macrophages, CSC-like cells, and cytotoxic T cells within CIC neighbourhoods (Figure 5B), indicating that these regions represent spatially distinct tumour microenvironments.

To further assess spatial organisation, we computed co-occurrence and neighbourhood enrichment scores. As expected, cancer cells showed the strongest co-occurrence with CICs. Notably, CSC-like cells were the only additional population exhibiting high co-occurrence and nearest-neighbour enrichment scores, indicating non-random spatial association with CICs (Figure 5C).

We next evaluated the clinical relevance of CIC neighbourhood composition. We grouped patients based on the presence or absence of cell types enriched in CIC neighbourhoods. Cancer or stromal cell presence within CIC neighbourhoods was not associated with clinical outcome (Figure 5D). Similarly, macrophages and CSC-like cells showed no association with survival. Notably, no deaths occurred among patients with cytotoxic T cells present within CIC neighbourhoods. When the analysis was extended to include all T cells to increase sample size, no deaths were observed in this group during follow-up (Figure 5E).

Because CICs exhibited increased glucose uptake, we hypothesised that they may deplete local glucose and influence cells within CIC neighbourhoods. We compared single-cell protein profiles of cytotoxic, helper, and regulatory T cells located within CIC neighbourhoods with those located more than 200 *µ*m away. Significant changes were observed only in cytotoxic T cells, which displayed increased GRP78, CASP8, CMYC, KI67, GLUT1, and β-catenin compared with distal cytotoxic T cells (Figure 5F).

We then analysed protein expression in cytotoxic T, helper T, regulatory T, cancer, and CSC-like cells as a function of distance from CICs (300 *µ*m to near proximity). GRP78 and GLUT1 expression increased progressively with proximity to CICs across multiple cell types, suggesting a distance-dependent spatial gradient (Figure 6). The largest increases were observed in cytotoxic T cells and CSC-like cells, beginning at approximately 180 *µ*m and 100 *µ*m from CICs, respectively, whereas other cell types showed more modest changes (Figure 6A–E). Additional glycolytic markers exhibited similar spatial trends, except for PKM2 in cytotoxic T cells and G6PD across cell types. CL-CASP3 levels remained unchanged in cytotoxic T cells, while these cells showed elevated CMYC and KI67 expression. In CSC-like cells, glycolytic proteins increased with proximity to CICs together with reduced BAK and increased BAX and CL-CASP3 expression.

**Figure 6:**
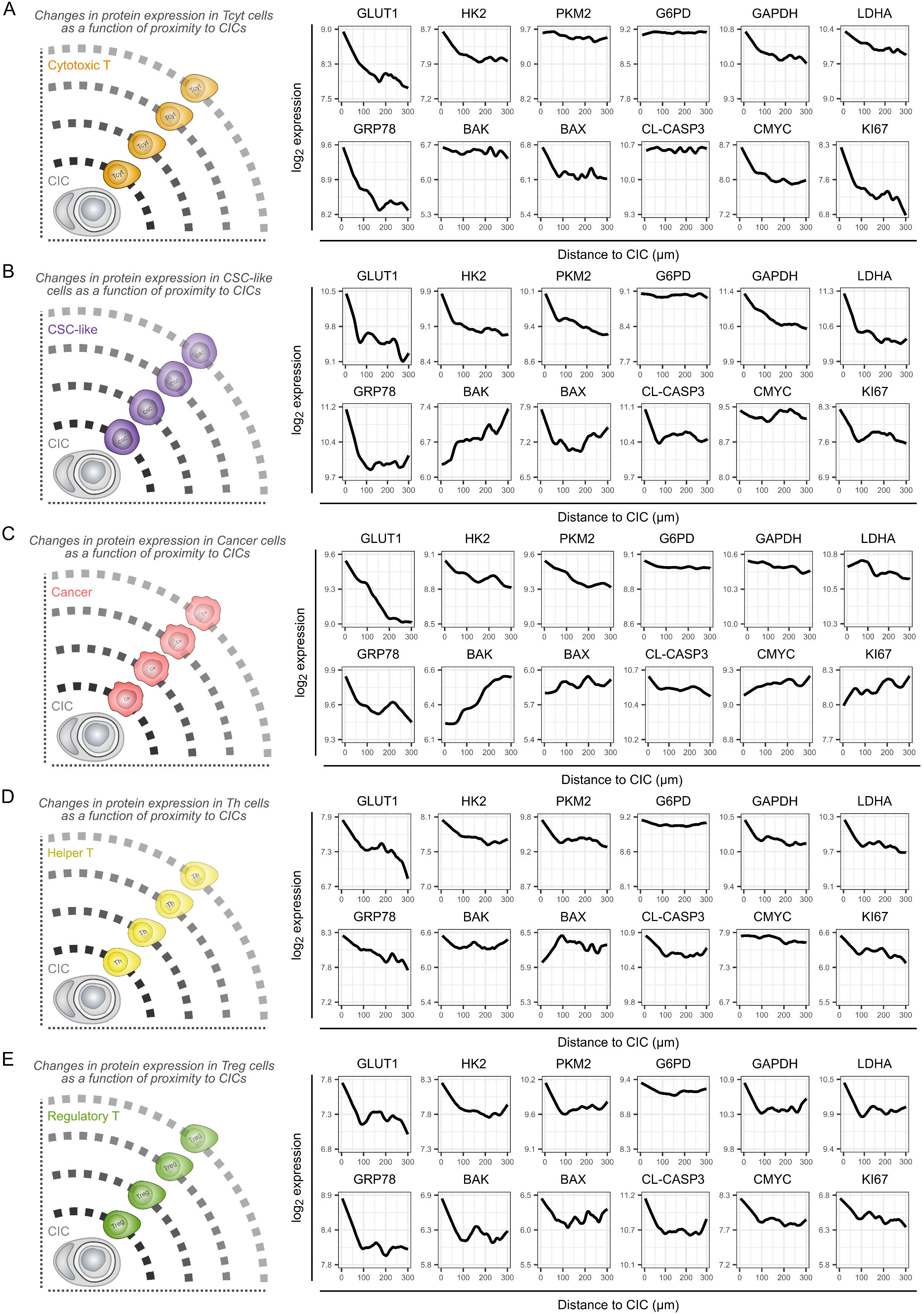
Proximity to CIC structures correlates with metabolic stress. Changes in apoptotic, metabolic, and proliferation markers in cytotoxic T cells (**A**) (Tcyt, n = 2902), CSC-like cells (**B**) (n = 3705), cancer cells (**C**) (n = 104543), helper T cells (**D**) (n = 3569), and regulatory T cells (**E**) (n = 3210) based on their proximity to a CIC event. Expression values were log_2_ normalised. Protein expression curves represent lowess fits computed across all analysed cells within each cell type.

Together, these findings indicate that CICs represent spatially distinct tumour microenvironments enriched in cancer and CSC-like cells. In a subset of patients, cytotoxic T cells were also present within CIC neighbourhoods and their presence was associated with improved long-term survival. Distance-dependent increases in metabolic stress markers, including GLUT1 and GRP78, suggest that CICs may function as local metabolic hubs that influence neighbouring cell states. CSC-like cells located near CICs also exhibited increased CL-CASP3 expression, consistent with a loser-cell phenotype. These findings suggest that cells residing within CIC neighbourhoods undergo distinct metabolic and apoptotic alterations that may influence their survival, functional state, and turnover in these regions.

### Spatial co-occurrence of CICs and CSC-like cells is an independent predictor of improved clinical outcome

Although cancer cells and CSC-like cells frequently co-existed with CICs, their presence within CIC neighbourhoods alone was not associated with patient outcome. To determine whether spatial interactions within CIC neighbourhoods showed prognostic significance, we first constructed cell-cell interaction networks based on single-cell spatial coordinates (Figure 7A). Closeness centrality and co-occurrence scores were calculated at the patient level for interactions between CICs and cancer cells, cytotoxic T cells, and CSC-like cells, and their associations with clinical outcomes were assessed using Cox proportional hazard models.

**Figure 7:**
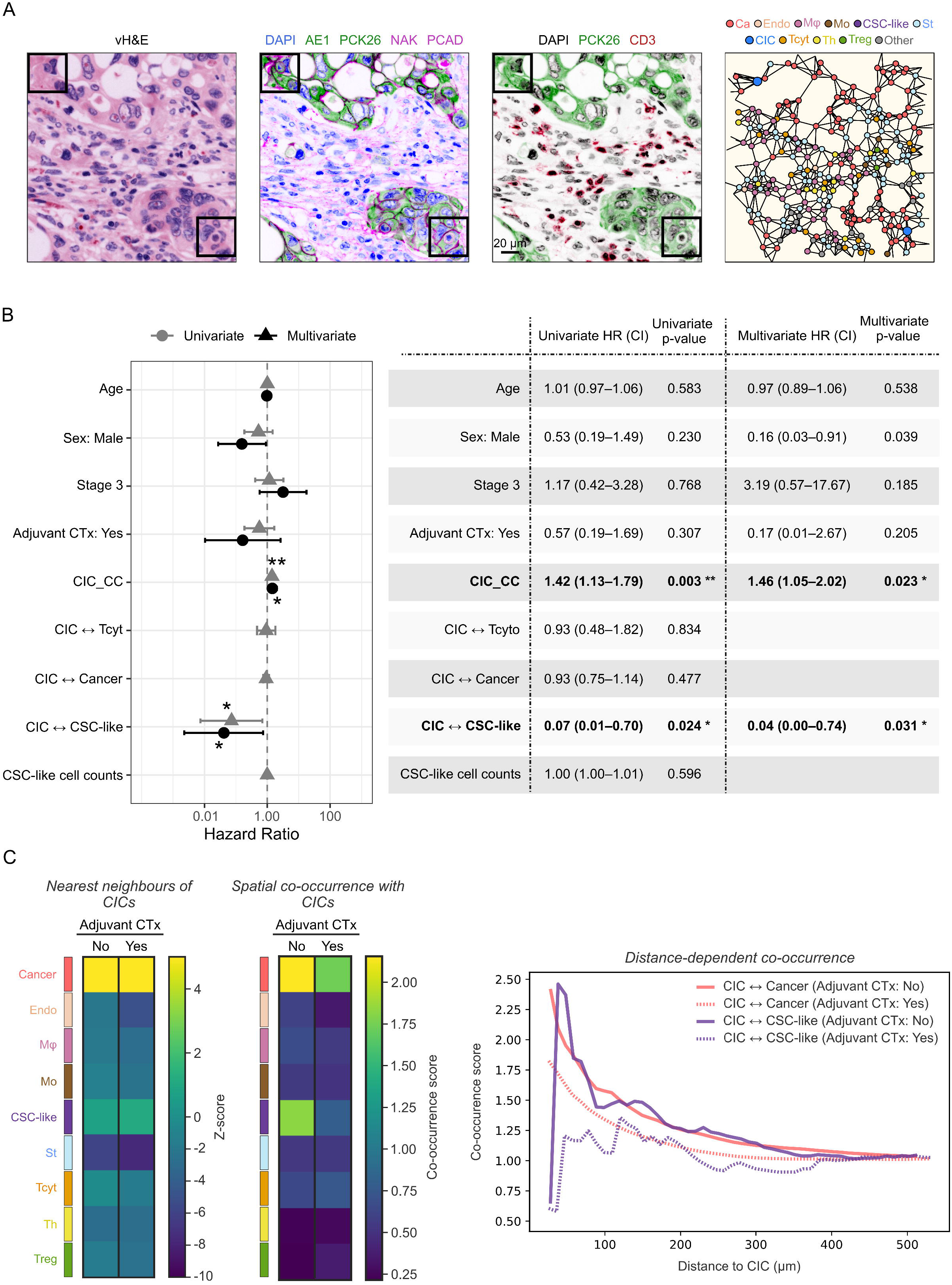
Association between spatial interactions within CIC neighbourhoods and clinical outcomes in colorectal cancer. **A**. Representative colorectal cancer tissue section and region of interest containing CICs (black squares) showing vH&E, staining of DAPI, AE1, PCK26, NAK, PCAD, CD3 along with the corresponding cell-cell interaction network map derived from spatial analysis. **B**. Univariate and multivariate analyses of spatial interactions within CIC neighbourhoods and age, sex, stage, adjuvant chemotherapy, and CSC-like cell counts in CRC patients (n = 94). To evaluate the prognostic significance of these features, Cox proportional hazard model was utilized to assess their impact on disease-free survival. Hazard ratios with 95% confidence intervals are shown. *P < 0.05, **P < 0.01. **C**. Adjuvant chemotherapy remodels spatial interactions between CICs and CSC-like cells. Co-occurrence probability and nearest-neighbour enrichment scores were compared in patients with (n = 69) or without (n = 78) adjuvant chemotherapy. Co-occurrence scores between CICs and cancer cells or CSC-like cells were plotted as a function of distance from CICs in patients with or without adjuvant chemotherapy.

Higher closeness centrality of CICs was significantly associated with poor patient outcome independent of age, sex, and stage [HR = 1.46 (1.05 - 2.02), p = 0.023]. In contrast, increased co-occurrence between CICs and CSC-like cells was associated with improved clinical outcome [(HR = 0.04 (0.00 - 0.74), p = 0.031)], and showed stronger prognostic value than age, sex, stage or adjuvant chemotherapy in this cohort. Co-occurrence between CICs and cytotoxic T cells or cancer cells, as well as the absolute number of CSC-like cells, were not associated with patient outcome, indicating that the effect was specific to spatial co-occurrence between CICs and CSC-like cells (Figure 7B).

We next examined the impact of adjuvant chemotherapy on CIC neighbourhood organisation by comparing nearest-neighbourhood enrichment and co-occurrence scores between CICs and each cell type. Adjuvant chemotherapy was associated with reduced co-occurrence between CICs and CSC-like cells at close proximity, whereas a more modest reduction was observed between CICs and cancer cells (Figure 7C). In contrast, adjuvant chemotherapy had minimal impact on the nearest-neighbour composition of CICs. These results reflect the distinction between the two metrics. Co-occurrence captures the overall frequency of CSC-like cells near CICs across the tissue, while nearest-neighbour enrichment measures immediate adjacency, which may remain stable even when CSC-like cells are globally less frequent. These results suggest that adjuvant chemotherapy may remodel CIC neighbourhoods by disrupting spatial interactions between CICs and CSC-like cells.

## Discussion

Cell competition has been extensively studied during development; however, its spatial organisation and clinical relevance in human tumours remain poorly understood. During cell competition, winner cells eliminate less fit neighbours through direct cell–cell interactions occurring at close proximity. One common mechanism involves induction of apoptosis^9^, while another involves cell extrusion^45^, in which neighbouring cells contract to expel the loser cell from the epithelial layer. In some contexts, winner cells eliminate competitors through engulfment, whereby outer cells internalise neighbouring loser cells and degrade them via lysosomal pathways^3,6,17^. In tumour tissues, this engulfment process has been reported for over a century as cell-in-cell structures (CIC)^15^. However, whether CICs represent competitive intratumoural microenvironments has remained unclear, largely due to the lack of approaches capable of resolving complex tissue architectures in tumours. Here, we systematically characterised the cellular identity and functional states of CICs *in situ*, examined their spatial interactions within the tumour microenvironment, and assessed their clinical relevance using single-cell, spatially resolved data from a large cohort of colorectal cancer patients. Our findings indicate that cell engulfment events represent spatially organised intratumoural niches, which we termed “CIC neighbourhoods”, that display molecular and spatial features consistent with cell competition and predict patient outcomes in colorectal cancer.

Consistent with previous studies^39,40^, CICs in colorectal tumours predominantly occur between cancer cells. Notably, we identified a previously unrecognised interaction in which ∼10% of CICs involved cancer stem cell (CSC)-like cells. Engulfed CSC-like cells exhibited increased CL-CASP3 and reduced CMYC expression, consistent with a loser-cell phenotype. Increased CL-CASP3 in CSC-like cells was also observed in cells located near CIC events, suggesting spatial interactions between these populations. Furthermore, spatial co-occurrence between CICs and CSC-like cells was associated with improved clinical outcomes, independent of age, sex, stage, and chemotherapy status. Interestingly, these spatial interactions were modified by adjuvant chemotherapy, suggesting that treatment may remodel competitive interactions within the tumour microenvironment. Given that CSCs constitute approximately 1.7%^46^ of colorectal cancer cells, engulfment of CSC-like cells by neighbouring cancer cells may represent a competitive mechanism similar to those described in normal intestinal epithelium^2,9^. Such interactions could influence tumour evolution by selectively eliminating particular cellular populations. *In vitro* models enabling inducible interactions between cancer cells and CSCs, such as intestinal crypt organoids, are required to investigate the mechanisms and consequences of cancer–CSC competition in CRC progression.

Nutrient availability within the tumour microenvironment generates heterogeneity in cellular metabolic fitness, which can drive supercompetition^2^. This is particularly relevant for cancer cells, as enhanced glycolysis even in the presence of oxygen, known as the Warburg effect, is a hallmark of cancer^47^. Cancer cells therefore produce large amounts of lactate, which acidifies the microenvironment and introduces additional metabolic competition, as lactate can also be used as an alternative energy source^48,49^. *In vitro* experiments show that glucose withdrawal promotes cell engulfment by generating metabolic heterogeneity between cancer cells through AMPK activity. Transient increase in AMPK activity is sufficient to define loser status in cells, which are then engulfed and eliminated by the winners. As a consequence, this process supports proliferation of winner cells under limited nutrient availability^24^. In parallel with previous work^23,24^, engulfment events between cancer cells were associated with upregulated glycolysis, particularly elevated GLUT1. Our observations are consistent with a model in which metabolic heterogeneity contributes to cell engulfment through competitive interactions. It will therefore be important to determine whether GLUT1 expression contributes to defining winner or loser cell status during competition, and whether it functions as driver or a consequence of CIC formation. In separate experiments, cancer cells exhibited increased glucose accumulation during degradation of inner cells, suggesting that cell engulfment events may locally deplete glucose. Spatial analysis revealed that cytotoxic T cells located near CICs exhibited metabolic stress markers associated with glucose deprivation, including Glucose-Regulated Protein 78 (GRP78)^50–54^, suggesting that cell engulfment may locally shape T cell functions. Indeed, competition for glucose between cancer cells and cytotoxic T cells within the tumour niches has been shown to suppress T cell effector functions^55,56^. In our study, cytotoxic T cells were also occasionally engulfed by cancer cells, and displayed elevated GRP78 and LDHA expression. Together with increased β-catenin levels in cytotoxic T cells located within CIC neighbourhoods, these findings suggest interactions consistent with enclysis^33^ and emperipolesis^31,32^, in which NK cells or cytotoxic T cells are engulfed and degraded by epithelial or tumour cells. Notably, no deaths were observed in patients with cytotoxic T cells present within CIC neighbourhoods, suggesting that these interactions may have clinical relevance.

Interestingly, both cancer and CSC-like cells associated with CICs exhibited lower expression of BCL-2 family proteins, particularly reduced BAK levels in inner cells. In addition to their well-established role in apoptosis^57^, experimental evidence from our laboratory^16^ and others^17,44^ indicates that deletion of *BAK* and *BAX* or overexpression of *BCL-2* can promote survival of inner cancer cells and delay or enable escape from degradation. These findings are also consistent with studies demonstrating that BCL-2 family proteins regulate cellular fitness during competitive interactions^5,6,13,14^. It will therefore be important to determine whether perturbations in BCL-2 family proteins influence competitive fitness in cancer cells. Given that cancer cells can initiate CIC formation in response to T cell attack^29^ or immune-associated cytokines such as TRAIL^16^, our findings also raise the possibility that dysregulation of BCL-2 family proteins may enable cancer cells to exploit CIC formation as a mechanism of immune evasion.

While this study provides novel insights into the spatial, functional, and clinical implications of cell engulfment in colorectal cancer, several limitations should be acknowledged. Because most analyses were performed on fixed tissue sections at a single time point, the temporal dynamics of cell competition and engulfment events cannot be fully resolved. Although distance-dependent spatial analyses partially mitigate this limitation, further *in vitro* studies are required to determine whether the protein expression changes observed in this study reflect underlying differences in metabolic activity. Whether CICs eliminate CSC-like cells through cell competition, suppress T cell infiltration, or arise as a consequence of pre-existing immune exclusion remains to be determined. Additionally, limited sample sizes may reduce statistical power in certain analyses, including protein expression and survival associations. Finally, although multiplex imaging enabled comprehensive spatial and phenotypic characterisation, rare instances of signal overlap or nonspecific staining cannot be completely excluded. These limitations highlight opportunities for further studies to clarify the mechanistic role of cell engulfment in cell competition, T cell exclusion, immune interactions, and tumour progression in colorectal cancer.

## Methods

### Patient Cohort

This study was conducted in accordance with ethical guidelines for clinical research and approved by Beaumont Hospital Research and Ethics Committee (References 08/62 and 19/46). Written informed consent was obtained from all participants. Clinical, demographic, and pathological characteristics of the patients are summarised in Table S2. Tissue microarrays (TMA) were constructed using three 1-mm-diameter cores from the tumour centre of formalin-fixed, paraffin-embedded (FFPE) primary tumour tissue sections for each of the 148 colorectal cancer patients (n = 444 cores).

### Multiplexed immunofluorescence (MxIF) imaging of TMAs

MxIF staining of the CRC TMAs was performed as previously described^58^ using Cell DIVE technology (Leica Microsystems, Issaquah, WA, USA), which allows for >60 protein markers to be stained, imaged and quantified at cell level within the same tissue section. Briefly, after deparaffinisation and two-step antigen-retrieval, TMAs were stained for 1 hour at room temperature using a Leica Bond autostainer. All antibodies were characterised and each antibody was conjugated with either Cy3 or Cy5 as described in the previous protocol^58^. TMAs were stained with DAPI and imaged every round in all channels of interest to acquire background autofluorescence. This was followed by antibody staining of up to three markers and DAPI each round, dye deactivation, and repeat staining to collect images of multiple biomarkers. All samples underwent autofluorescence background imaging for the first five rounds and every three rounds thereafter. Markers of interest and corresponding antibodies used in this study are listed in Table S1.

### Image processing

Background autofluorescence subtraction, illumination and distortion correction were performed using Cell DIVE automated image pre-processing software. DAPI and Cy3 autofluorescence images were used to generate virtual H&E (vH&E) images. Cells in the epithelial and stromal compartments were segmented using DAPI, pan-cytokeratin, S6 and NaKATPase, and each segmented cell was assigned a unique ID and spatial coordinate, as previously described^58^. After segmentation, several quality control (QC) steps were conducted^59,60^. Briefly, each image was reviewed for tissue completeness and accuracy of segmentation masks in each subcellular compartment as well as epithelial and stromal tissue separation. Images with poor staining or poor segmentation were excluded from data analysis. The following criteria were applied for cell filtering: (1) epithelial cells to have either one or two nuclei; (2) area of each subcellular compartment (nucleus, membrane, cytoplasm) to be >10 pixels and <1,500 pixels; (3) cell to show excellent alignment with the first round of staining; (4) cells to be located >25 pixels distance from the image margins; (5) cell area for nuclear segmentation mask to be >100 and <3,000 pixels, (6) duplicates. After comprehensive QC checks, the data were further processed and normalised to remove batch effects, and log2 transformation was performed to handle the skewness of the marker intensities as previously described^60,61^.

### Cell type characterisation

Cell types were characterised by the classification pipeline developed in our lab and described in^62^. Briefly, we classified CDX2+ PCK26+ cells as cancer, CDX2-PCK26-AE1-cells as stromal, CD31high COLIVhigh SMAhigh cells as endothelial, CD4high CD68high cells as macrophages, CD3-CD8-CD4+ cells as monocytes, ALDHhigh KI67low AE1low cells as cancer stem cell-like (CSC-like), CD3+ CD8+ cells as cytotoxic T, CD3+ CD4+ cells as helper T, and CD3+ CD4+ FOXP3+ cells as regulatory T cells. In total, 743429 cells were characterised across 344 cores from 124 patients.

### Characterisation of cell-in-cell events

Cell-in-cell structures were identified using images of virtual H&E and a combination of nuclear (DAPI), cytoplasmic (AE1, PCK26, EPCAM), and membrane (NAK, PCAD) markers. CICs were manually annotated using HALO software if they exhibited evidence of a complete internalisation of an entire cell within another cell and if the events demonstrated at least five of the following six criteria: (1) nucleus of inner cell, (2) cytoplasm of inner cell, (3) membrane of inner cell, (4) nucleus of outer cell, (5) cytoplasm of outer cell, and (6) visible vacuolar space between inner and outer cells, similar to previous studies^16,20,43^.

### Spatial analysis

Proximity and network analyses were performed as previously described^63,64^. Briefly, for the proximity analysis, Euclidean distances between the centre points of cell types were calculated for each cell, and the average distances were used to compare TMA cores. A 20 *µ*m distance threshold was used to define CIC neighbourhoods, consistent with previous studies that considered cells within this range as adjacent^65^. For the network analysis, the Squidpy^66^ library was used to create network graphs based on cell coordinates. Closeness centrality, co-occurrence and neighbourhood enrichment scores were calculated for each tumour core separately and averaged for patient-level comparisons. Co-occurrence scores aggregated within 100 *µ*m were used for clinical comparisons.

### Cell culture and reagents

HCT116 and Jurkat cells were cultured in RPMI medium supplemented with 10% FBS, 2 mM L-glutamine, 100 U/ml penicillin, and 100 μg/ml streptomycin at 37^°^C in a humidified atmosphere at 5% CO_2_. Cell lines were authenticated at the beginning and completion of the study and regularly tested for mycoplasma. Dimethyl sulfoxide (#D8418), Dulbecco’s PBS (#D8537), FBS (#F7525), Hanks’ balanced salt solution (#H8264), Hoechst 33258 (#94403), L-glutamine (#G7513), penicillin-streptomycin (#P0781), Roswell Park Memorial Institute 1640 medium (RPMI medium; #R0883), and trypsin-EDTA (#T4049) were purchased from Sigma-Aldrich. SILAC RPMI medium (#A24942-01, Gibco), dialyzed FBS (dFBS, #A33820-01, Gibco), LysoTracker Deep Red (#L12492), CellTracker Deep Red (#C34565), 2-NBDG (2-(N-(7-Nitrobenz-2-oxa-1,3-diazol-4-yl)Amino)-2-Deoxyglucose) (#N13195) were obtained from Thermo Fisher Scientific.

### Microscopy

#### Live-cell confocal microscopy imaging of glucose (2-NBDG) uptake

Cells were seeded on sterile 12-mm glass-bottom WillCo-dishes (WillCo Wells B.V.) and allowed to adhere overnight in RPMI medium at 37°C with 5% CO_2_. Next day, cells were incubated in staining medium containing a 1 *µ*g/ml Hoechst 33342, 100 *µ*M 2-NBDG, and 5 nM LysoTracker Deep Red. WillCo-dishes containing stained cells were covered with embryo-tested sterile mineral oil and mounted on a LSM 980 Airyscan 2 Confocal Microscope equipped with an incubation chamber (Carl Zeiss). Images were acquired for 24 hours with 5-minute intervals. For co-culture experiments (Figure S3A), Jurkat cells were pre-stained with 2 *µ*M CellTracker Deep Red for 15 min in prewarmed RPMI medium, washed three times with PBS, and co-cultured with HCT116 at a 1:1 ratio. Co-cultures were incubated in staining medium containing 1 *µ*g/ml Hoechst 33342, 100 *µ*M 2-NBDG, and 200 nM Lysotracker Red for 12 hours. Imaging was performed using SP8 confocal laser-scanning microscope equipped with an incubation chamber (Leica). Images were analysed using ImageJ (v1.54p)^67^ as previously described^27,68^.

#### Immunofluorescence staining and 3D confocal imaging

Cells were seeded on sterile 12-mm glass-bottom WillCo-dishes (WillCo Wells B.V.) and allowed to adhere overnight in RPMI or SILAC medium (with dFBS) at 37°C with 5% CO_2_. Next day, cells were incubated in staining medium containing a 1 *µ*g/ml Hoechst 33342 and 5 nM LysoTracker Deep Red for 24 hours. After incubation, cells were fixed with 4% paraformaldehyde in PBS for 20 minutes at room temperature, washed three times with PBS, and blocked for 1 hour. Cells were then incubated with Alexa Fluor 488-conjugated anti-GLUT1 antibody (#07-1401-AF488, Sigma-Aldrich) for 2 hours, washed three times with PBS, and imaged using LSM 980 Airyscan 2 Confocal Microscope (Carl Zeiss). Images were analysed using ImageJ (v1.54p)^67^.

### Statistical analysis

All statistical tests were performed using R (v4.4.3; R Core Team 2025, RRID:SCR_001905) and P values < 0.05 were considered statistically significant. Data are presented as median with interquartile range, and mean. Two-tailed t-tests were used for pairwise comparisons between two groups. For comparisons involving more than two groups, one-way ANOVA followed by Tukey’s Honestly Significant Difference (HSD) post-hoc test was performed. Differences in survival curves between patient groups were assessed by Kaplan–Meier estimates and log-rank tests. Associations with survival were further analysed using multivariable Cox proportional hazard models adjusted for age and sex. P values, hazard ratios (HR), and 95% confidence intervals (CI) were reported. Power analysis was not performed, as this was an exploratory pilot study. No samples or data points were excluded unless otherwise mentioned. Sex was included as a biological variable in multivariable models where relevant. Patients were not randomised, as this study was observational and based on existing tissue collections. All CIC annotations and subsequent analyses were conducted blinded to tissue identity and clinical outcomes. Detailed descriptions of statistical tests and data presentations are provided in the figure legends. All experiments were repeated at least three times. Asterisks indicate significance as follows *P < 0.05, **P < 0.01, ***P < 0.001, ns: not significant.

## Acknowledgements

Research reported in this publication was supported by a grant from Research Ireland and the Health Research Board (18/RI/5792, 21/RI/9787, 16/US/3301 and ERA-TRANSCAN-2022-002) to JHMP. FG was supported by the National Cancer Institute of the National Institutes of Health (R01CA208179). DBL was supported by a US-Ireland R01 award (NI Partner supported by HSCNI, STL/5715/15). BK was supported by Research Ireland through Centre for Research Training in Genomics Data Science (18/CRT/6214), and the European Union Horizon 2020 research and innovation programme under the Marie Sklodowska-Curie grant (H2020-MSCA-COFUND-2019-945385). FL was funded by MICIU/AEI/10.13039/501100011033 (PID2023-152599OA-I00 and RYC2021-034092-I). We also thank all study participants for their contribution.

## Authors’ Contributions

Conceptualisation: EB, JHMP. Methodology: EB, BK, MA, HD, AS, FL, SC, EMcD, JF, TO’G. Formal analysis: EB, BK. Investigation: EB. Resources: JPB, NMcC, DAMcN. Writing–original draft: EB. Writing–review and editing: EB, JHMP. Visualization: EB, BK. Funding acquisition: JHMP, FG, DBL. All authors read and reviewed the manuscript.

## Authors’ Disclosures

No disclosures were reported by the authors.

## Data availability

Further information and requests for datasets, code, or additional resources related to this study will be fulfilled by the corresponding author upon request.

## Notes

### Competing Interest Statement

The authors have declared no competing interest.

### Summary of Updates

This manuscript has been substantially updated. The title, framing, author list, and content have been revised throughout, with updated figures and supplemental materials.

