## Supplementary figures and images for "Cell engulfment defines spatially distinct competitive metabolic niches associated with clinical outcomes in colorectal cancer"

### Supplementary Figure 1

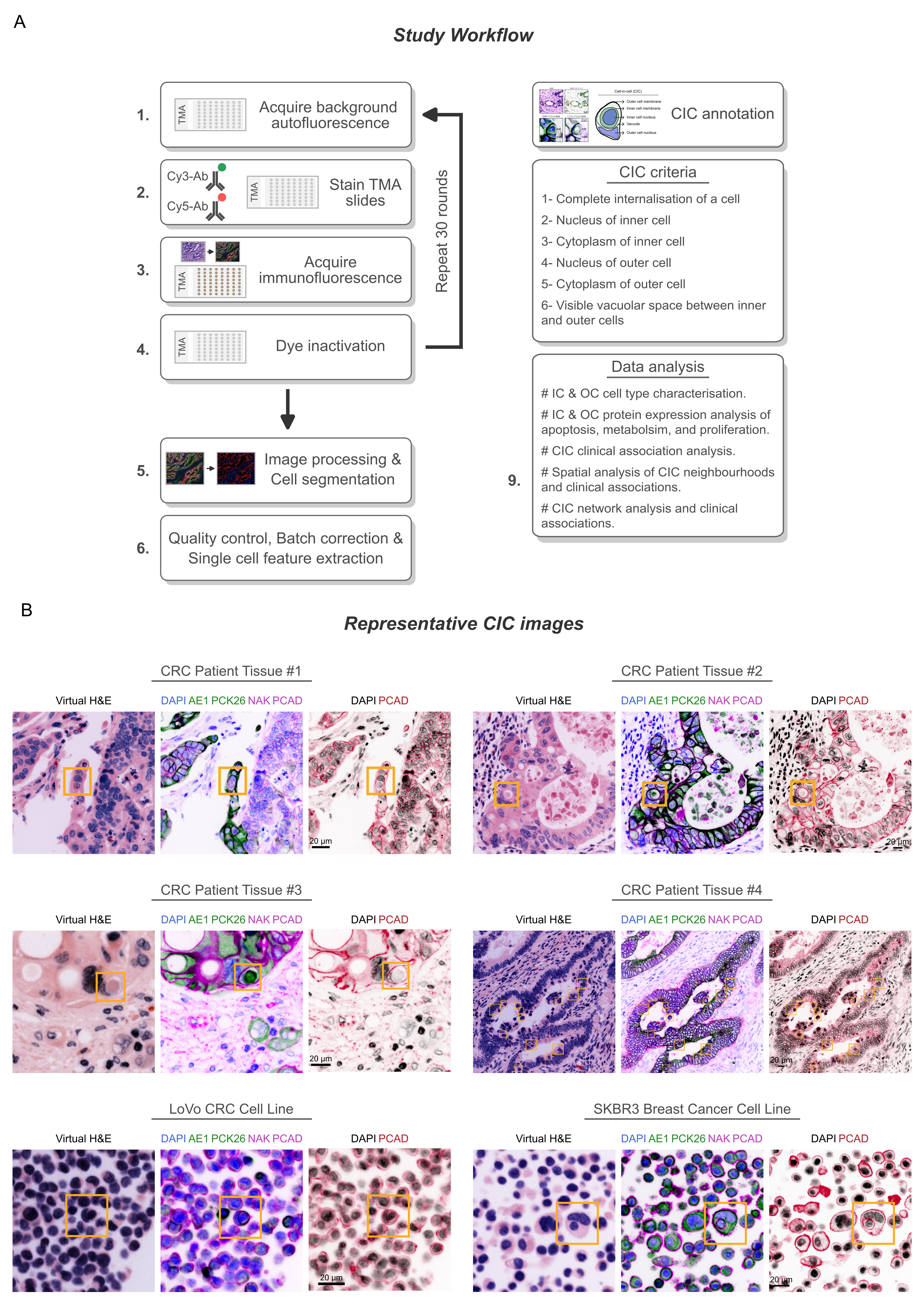

### Supplementary Figure 2

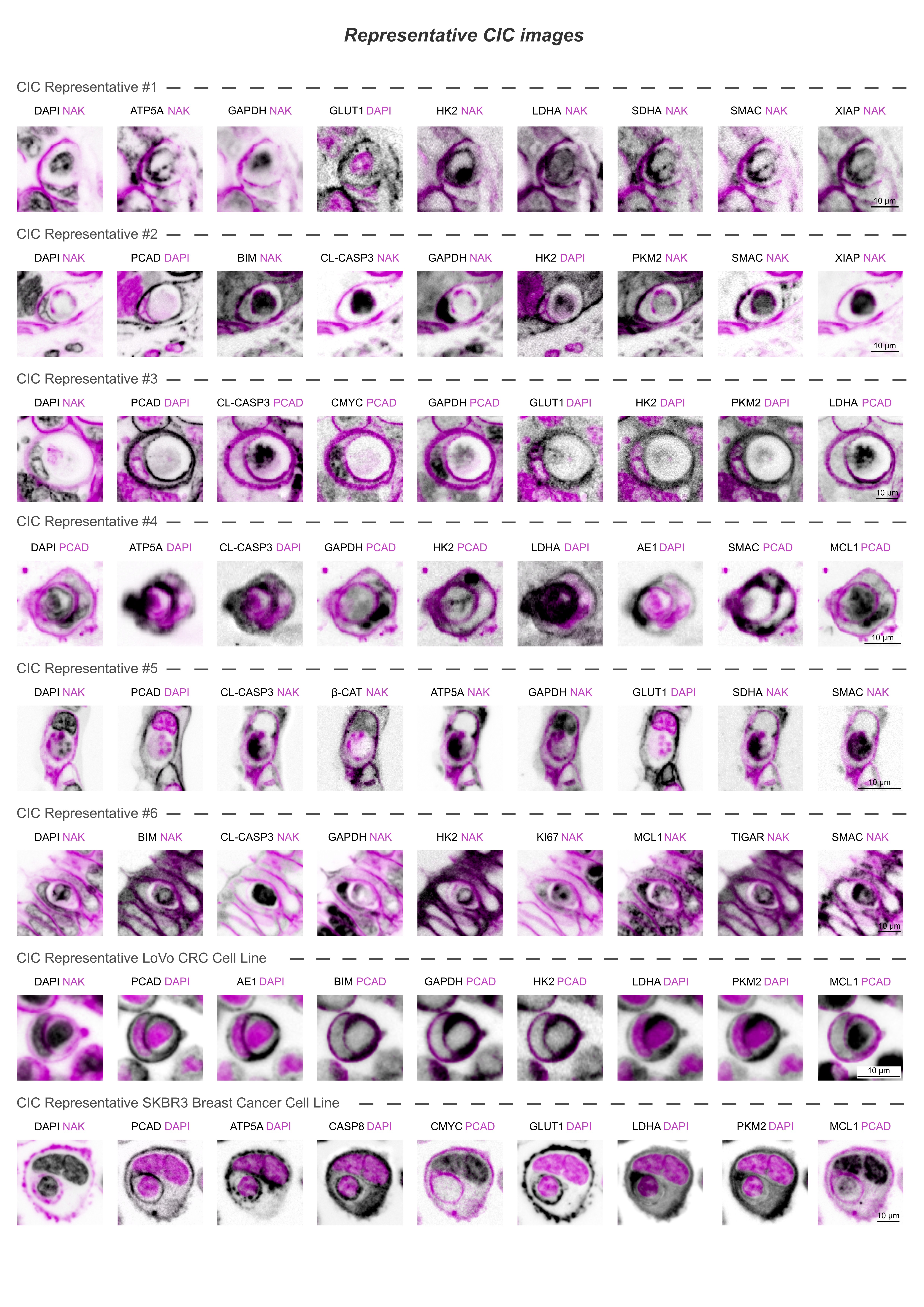

### Supplementary Figure 3

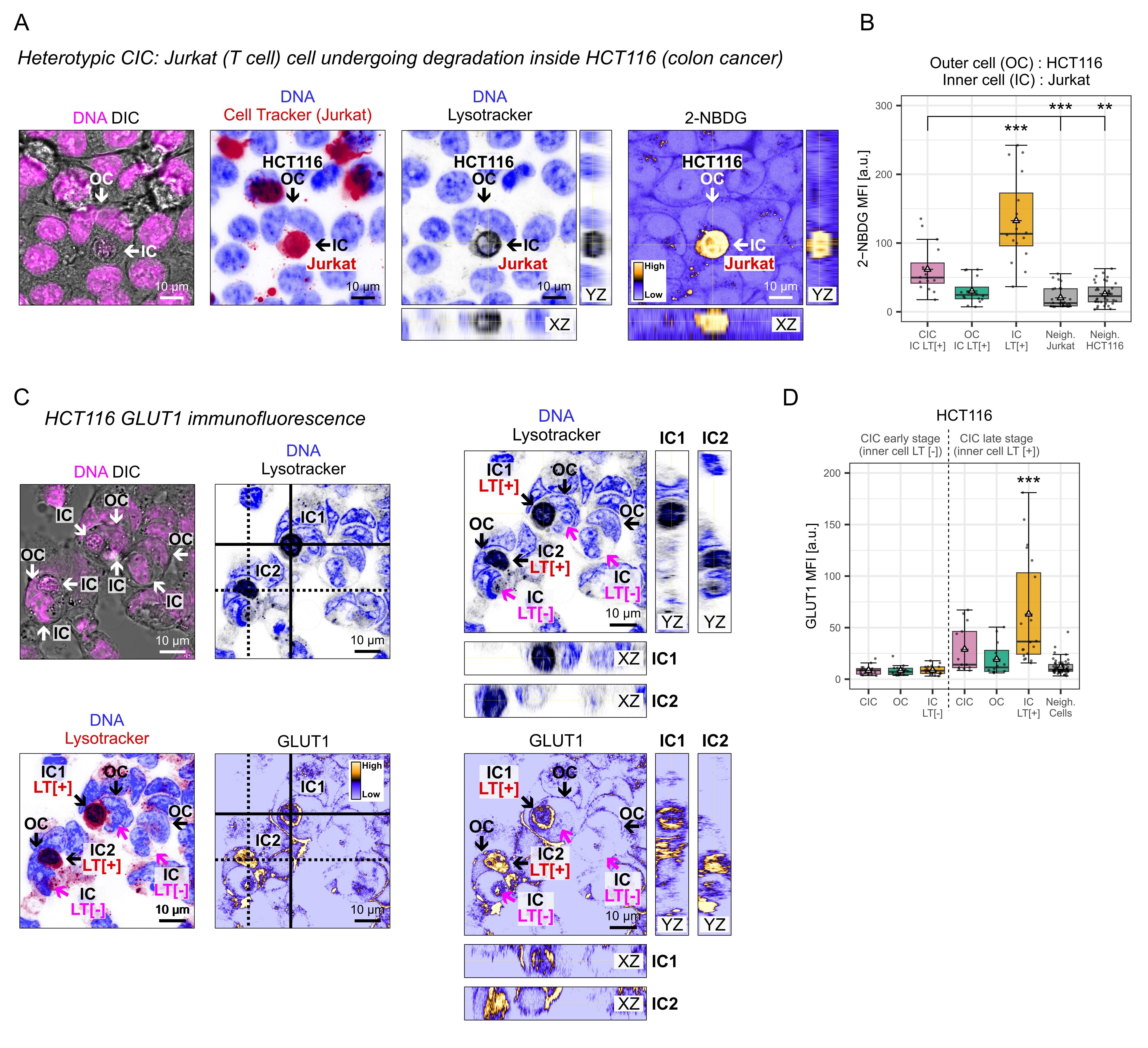
