## Supplementary Information for "Cell engulfment defines spatially distinct competitive metabolic niches associated with clinical outcomes in colorectal cancer"

**Supplementary Figure 1**: **A.** Simplified workflow of Cell DIVE multiplex imaging, CIC annotation and data analysis. **B.** Representative CRC tissue sections and cell lines showing vH&E and staining of DAPI, AE1, PCK26, NAK, and PCAD. Yellow squares highlight regions of interest containing cell-in-cell (CIC) events. Combination of vH&E, nuclear (DAPI), cytoplasmic (AE1, PCK26) and membrane (NAK, PCAD) markers was used to identify CICs.

**Supplementary Figure 2**: Representative images of CICs in CRC tissue sections and cell lines showing the expression profiles of proteins involved in key cellular processes, including apoptosis, metabolism, and proliferation in inner and outer cells. Nuclei (DAPI) or cell membrane (NAK or PCAD) are shown in purple.

**Supplementary Figure 3**: **A.** Representative images of DNA (Hoechst), Cell Tracker (Jurkat cells), Lysotracker, and orthogonal views of 2-NBDG staining showing a Jurkat cell (inner cell) undergoing lysosomal degradation within HCT116 cell (outer cell). **B.** Quantification of 2-NBDG mean fluorescence intensity (MFI) in heterotypic CICs containing HCT116 as outer cells (n = 16) and Jurkat cells as inner cells undergoing lysosomal degradation (ICLT[+], n = 16), as well as in neighbouring HCT116 (n = 36) and Jurkat cells (n = 26). Groups were compared using one-way ANOVA followed by Tukey’s HSD post-hoc test for multiple comparisons. **P < 0.01, ***P < 0.001. **C.** Representative images of DNA (Hoechst), Lysotracker, and GLUT1 immunofluorescence staining showing homotypic CICs between HCT116 cells. A field of view containing multiple CICs is shown, along with orthogonal views of DNA, Lysotracker, GLUT1 staining of two representative events (IC1, IC2). LT [+] (red) and LT [-] (magenta) inner cells are indicated. OC: outer cell, IC: inner cell. **D.** Quantification of GLUT1 mean fluorescence intensity (MFI) in homotypic CICs between HCT116 cells. Early-stage CICs contain inner cells without Lysotracker accumulation (LT [-]), whereas late-stage CICs contain inner cells undergoing lysosomal degradation (LT [+]). Numbers of analysed cells for each group were: nCIC_ICLT[-] = 17, nOC_ICLT[-] = 17, nICLT[-] = 19, nCIC_ICLT[+] = 13, nOC_ICLT[+] = 12, nICLT[+] = 20, Neigh. cells = 57. Groups were compared using one-way ANOVA followed by Tukey’s HSD post-hoc test for multiple comparisons. ***P < 0.001.

**Supplementary Table 1**: List of proteins, antibodies, manufacturers, catalogue numbers, and concentrations used in this study.

**Supplementary Table 2**: Associations between the absence or presence of CICs and the clinical, demographic, and pathological characteristics of the patient cohort. Continuous variables are summarized with mean ± SD, median (Q1–Q3), and range. Categorical variables are presented as counts and row percentages. N-miss indicates missing data.
